# The Uvrag-containing PI3K complex promotes Hsc70-4 dependent endosomal clathrin removal and lysosomal maturation in Drosophila nephrocytes

**DOI:** 10.64898/2026.05.12.724521

**Authors:** Anikó Nagy, Virág Balogh, Dávid Hargitai, Attila Boda, Enikő Horváth, Zsófia Simon-Vecsei, Gábor Juhász, Péter Lőrincz

## Abstract

The class III phosphatidylinositol 3-kinase complex (PI3K(III)) generates phosphatidylinositol-3-phosphate (PI(3)P), a lipid that defines endosomal membrane identity. Two PI3K(III) complexes share core subunits but differ in their fourth component: the Atg14-containing complex I functions in autophagy, whereas the Uvrag-containing complex II is required for endosomal maturation. Despite this, the mechanism by which complex II promotes lysosomal function remains unclear. Using Drosophila nephrocytes, we show that PI(3)P is enriched on Rab7-positive late endosomes and that the Hsp70 chaperone Hsc70-4 binds phosphoinositides. Loss of PI3K complex II disrupts endolysosomal organization and phenocopies Hsc70-4 inhibition. In both cases, clathrin accumulates on intracellular, often endosomal membranes, Rab7 compartments are disorganized, and abnormal endolysosomal structures form. These defects are accompanied by impaired HOPS recruitment, lysosomal dysfunction, and secretion of endolysosomal content. Importantly, clathrin depletion partially rescues these defects. Together, our findings identify a role for PI3K complex II in promoting clathrin removal from endosomal membranes and link PI(3)P and Hsc70-4 activity to lysosomal maturation.

## Introduction

Endocytic trafficking relies on a highly coordinated maturation process in which newly formed vesicles undergo sequential changes in protein and lipid composition. Central to this process are Rab small GTPases, which define the identity of endosomal compartments and regulate vesicle movement, tethering, and fusion. Rab proteins recruit specific effector complexes, including multisubunit tethering complexes such as the homotypic fusion and vacuole protein sorting (HOPS) complex, which promote membrane tethering and facilitate SNARE-mediated endosome–lysosome fusion. Through these mechanisms, Rab proteins and their effectors drive the orderly progression of cargo from early endosomes toward late endosomes and lysosomes (Wickner and Schekman 2008, Doherty and McMahon 2009).

Membrane identity during endosomal maturation is also strongly influenced by phosphoinositide lipids (Falkenburger, Jensen et al. 2010). The class III phosphatidylinositol 3-kinase (PI3K(III)) complex plays a central role in this process by generating phosphatidylinositol-3-phosphate (PI(3)P) from phosphatidylinositol on specific intracellular membranes. PI(3)P is enriched on early endosomes, autophagosomes, and multivesicular bodies and serves as a docking platform for proteins containing for example FYVE or Phox homology (PX) domains that bind to this phospholipid, thereby coordinating downstream membrane trafficking events (Gillooly, Morrow et al. 2000, Gillooly, Simonsen and Stenmark 2001, Backer 2008).

In Drosophila melanogaster, two PI3K(III) complexes share three core subunits: the lipid kinase Vps34, the regulatory kinase Vps15, and the scaffold protein Atg6 (Vps30 in yeast and Beclin1 in mammals). Vps34 provides the catalytic activity responsible for PI(3)P production, while Vps15 and Atg6 regulate complex stability, localization, and activity. The functional specificity of the complexes is determined by their fourth subunit: complex I contains Atg14 and functions primarily in autophagosome biogenesis, whereas complex II contains Uvrag (Vps38 in yeast) and is required for endosome maturation and lysosomal trafficking (Kihara, Noda et al. 2001, Backer 2008, Itakura, Kishi et al. 2008, Funderburk, Wang and Yue 2010).

Multiple studies, including our previous work, have shown that loss of complex II function disrupts lysosomal maturation (Juhász, Hill et al. 2008, Abe, Setoguchi et al. 2009, Lee, Liang et al. 2011, Lorincz, Lakatos et al. 2014, Takats, Pircs et al. 2014, Munson, Allen et al. 2015). In these conditions, abnormal endolysosomal compartments accumulate and display impaired degradative capacity. Although these findings demonstrate the requirement of complex II for lysosomal function, the underlying mechanism remains unclear.

Hsc70 proteins belong to the Hsp70 family of molecular chaperones and participate in diverse cellular processes, including protein folding, membrane remodeling, and vesicle trafficking (McMahon and Boucrot 2011). The Drosophila homolog of mammalian HSPA8/Hsc70 is Hsc70-4. In neurons, Hsc70-4 promotes endosomal microautophagy (eMI), during which endosomes internalize cytosolic cargo and mature into multivesicular bodies (Uytterhoeven, Lauwers et al. 2015). Importantly, Hsc70-4 plays a well-established role in endocytosis by mediating the uncoating of clathrin-coated vesicles following internalization (DeLuca-Flaherty, McKay et al. 1990, Cheetham, Anderton and Jackson 1996, Newmyer and Schmid 2001, Chang, Newmyer et al. 2002, Eisenberg and Greene 2007, Böcking, Aguet et al. 2011).

The larval garland nephrocyte of Drosophila provides a powerful model to study endocytosis due to its exceptionally high endocytic activity. These cells display a highly organized endolysosomal architecture in which endocytic compartments form distinct cytoplasmic layers: clathrin-coated vesicles are located near the plasma membrane, followed by early and late endosomes, while lysosomes accumulate near the nucleus (Lorincz, Lakatos et al. 2016, Lorincz, Toth et al. 2017, Lorincz, Kenez et al. 2019). This spatial organization facilitates the analysis of defects in endocytic trafficking. In addition, garland nephrocytes possess a vertebrate-like slit diaphragm and are widely used as a model system for studying nephropathies (Simons and Huber 2009, Weavers, Prieto-Sanchez et al. 2009, Zhuang, Shao et al. 2009, Helmstädter, Huber and Hermle 2017). Notably, Hsc70-4 is expressed in these cells, and its role in clathrin uncoating has been demonstrated in nephrocytes, in part through its interaction with receptor-mediated endocytosis 8 (Rme-8) (Chang, Newmyer et al. 2002, Chang, Hull and Mellman 2004).

Here we report that loss of PI3K(III) complex II produces phenotypes strikingly similar to those observed upon Hsc70-4 inhibition. In these conditions, clathrin remains associated with endosomal membranes, suggesting defective clathrin uncoating that may interfere with endosomal maturation and lysosomal membrane recycling. As a consequence, lysosomal function is impaired, leading to lysosomal exocytosis and increased recycling of endosomal membranes. Furthermore, these abnormal compartments fail to recruit the HOPS tethering complex, indicating a defect in the acquisition of late endosomal identity. Our findings identify a previously unrecognized link between PI(3)P production, clathrin uncoating, and lysosomal maturation.

## Results

### PI3K(III) complex II is required for the organization of late endosomes and lysosomes in nephrocytes

To investigate the role of PI3K(III) complex II in endolysosomal organization, we analyzed Drosophila larval garland nephrocytes, a cell type with exceptionally high endocytic activity and a highly ordered endolysosomal architecture. In these cells, endocytic compartments are arranged in distinct cytoplasmic layers, with clathrin-coated vesicles located near the plasma membrane, followed by early endosomes, late endosomes, and lysosomes toward the cell interior (Lorincz, Lakatos et al. 2016). To monitor PI(3)P distribution and late endosome identity, nephrocytes expressing the PI(3)P reporter GFP-FYVE were immunostained for the late endosomal marker Rab7. In control cells, GFP-FYVE–positive structures were present in both peripheral early endosomes and deeper cytoplasmic regions, where they strongly colocalized with Rab7 on large late endosomes, indicating enrichment of PI(3)P (Fig. 1A,Q). A similar pattern was observed upon depletion of Atg14, a subunit specific for PI3K complex I (Fig. 1D,Q), suggesting that complex I is largely dispensable for maintaining PI(3)P-positive late endosomes in nephrocytes. In contrast, depletion of the catalytic subunit Vps34 or the complex II–specific subunit Uvrag resulted in a marked reduction of GFP-FYVE signal (Fig. 1B,C,Q). Under these conditions, Rab7-positive structures were strongly altered: the typical large late endosomes were absent, and Rab7 signal accumulated either at the cell periphery or in abnormal intracellular structures (Fig. 1B,C). Similar phenotypes were observed upon depletion of the regulatory PI3K complex subunits Vps15 and Atg6 (Supplementary Fig. 1A-C), indicating that PI3K complex activity is required for maintaining normal late endosomal organization.

**Figure 1.**
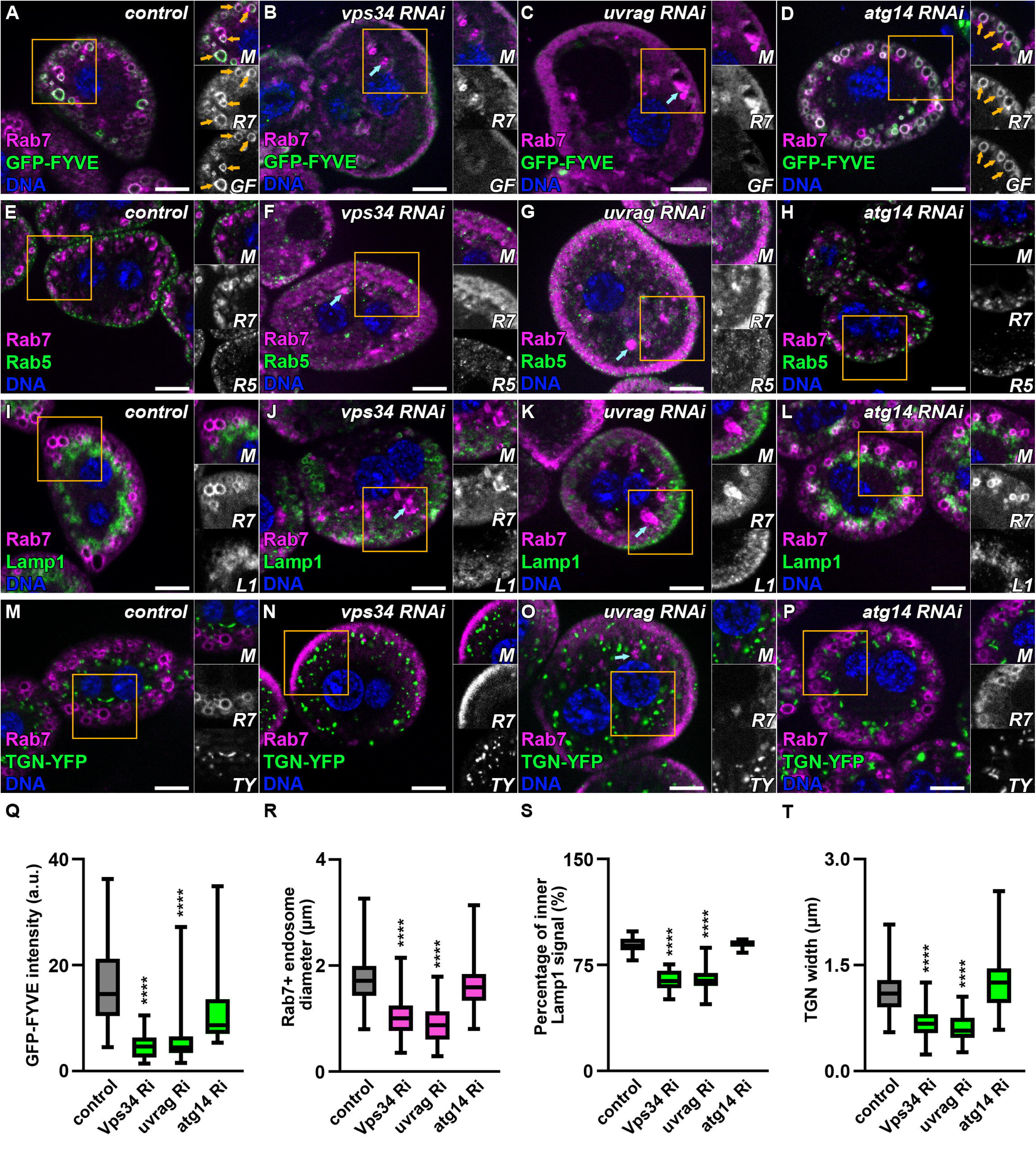
PI3K(III) complex II is required for normal late endosome and lysosome organization in nephrocytes. **A–D.** Large Rab7-positive late endosomes are observed in control nephrocytes (A) and in cells depleted of Atg14 (PI3K(III) complex I–specific subunit) (D), where Rab7 strongly colocalizes with the PI(3)P reporter GFP-FYVE (orange arrows). In contrast, depletion of the catalytic subunit Vps34 (B) or the complex II–specific subunit Uvrag (C) leads to a strong reduction of GFP-FYVE signal. Under these conditions, most Rab7 signal relocates to the cell periphery or forms large intracellular structures (blue arrows). Nuclei are shown in blue (DNA). Scale bars: 10 μm. Insets display merged (M), Rab7 (R7), and GFP-FYVE (GF) channels. **E–H.** Early endosomes were labeled using anti-Rab5 and late endosomes using anti-Rab7 antibodies. In control cells (E) and Atg14 knockdown cells (H), late endosomes form a distinct layer beneath the Rab5-positive early endosomes. In contrast, in Vps34- (F) or Uvrag-depleted (G) cells, Rab7-positive endosomes are largely lost, with remaining Rab7 signal accumulating at the cell periphery or forming large intracellular structures (blue arrows). In these cells, the Rab5 and Rab7 layers appear intermixed. Nuclei are shown in blue (DNA). Scale bars: 10 μm. Insets display merged (M), Rab5 (R5) and Rab7 (R7) signals of the boxed regions. **I–L.** Late endosomes were labeled with anti-Rab7 and lysosomes with anti-Lamp1. In control (I) and Atg14 knockdown cells (L), Lamp1-positive lysosomes form a layer near and slightly beneath Rab7-positive late endosomes. In contrast, in Vps34- (J) or Uvrag-depleted (K) cells, this organization becomes disrupted: both Rab7 and Lamp1 signals redistribute to the cell periphery or accumulate in large intracellular structures (blue arrows). Nuclei are shown in blue (DNA). Scale bars: 10 μm. Insets display merged (M), Rab7 (R7), and Lamp1 (L1) signals of the boxed regions. **M–P.** Nephrocytes expressing TGN-YFP were labeled with anti-Rab7. In control (M) and Atg14 knockdown cells (P), Golgi structures appear elongated and are located beneath late endosomes. In contrast, in Vps34- (N) or Uvrag-depleted (O) cells, TGN-YFP–positive structures appear more rounded and less organized. Nuclei are shown in blue (DNA). Scale bars: 10 μm. Insets display merged (M), Rab7 (R7), and TGN-YFP (TY) signals of the boxed regions. **Q.** Quantification of GFP-FYVE fluorescence intensity shown in panels A–D (n = 44). **R.** Quantification of Rab7-positive endosome size shown in panels E–H (n = 100). **S.** Quantification of the proportion of inner Lamp1 signal shown in panels I–L (n = 15). **T.** Quantification of the maximum width of Golgi-YFP structures shown in panels M–P (n = 100).

We next examined the spatial organization of endosomal compartments using Rab5 and Rab7 as markers of early and late endosomes, respectively. In control nephrocytes and Atg14-depleted cells, Rab5-positive early endosomes formed a peripheral layer, while Rab7-positive late endosomes localized in a distinct layer beneath them (Fig. 1E,H,R). However, this organization was disrupted in PI3K(III) complex II (Uvrag, Vps34, Vps15 or Atg6) depleted cells. In these cells, Rab7-positive compartments were again largely lost and the remaining Rab7 signal appeared either peripheral or in intracellular structures, while the Rab5 and Rab7 layers became intermixed (Fig. 1F,G, Supplementary Fig. 1D-N).

We further analyzed lysosomal organization by staining nephrocytes for Rab7 together with the lysosomal marker Lamp1. In control and Atg14 knockdown cells, Lamp1-positive lysosomes formed a layer near and slightly beneath Rab7-positive late endosomes (Fig. 1I,L). In contrast, depletion of PI3K(III) complex II proteins (Uvrag, Vps34, Vps15 or Atg6) disrupted this organization: both Rab7 and Lamp1 signals redistributed toward the cell periphery or accumulated in large intracellular structures (Fig. 1J,K,S, Supplementary Fig. 1O-Q).

Because lysosomal proteins are delivered from the Golgi apparatus, we next examined Golgi morphology using TGN-YFP. In control and Atg14-depleted cells, Golgi structures appeared elongated and were positioned beneath late endosomes (Fig. 1M,P,T). In contrast, in Vps34-or Uvrag-depleted cells, TGN-YFP–positive structures appeared more rounded and their specific distribution was lost (Fig. 1N,O,T), suggesting that PI3K complex II activity also contributes to proper Golgi organization.

Consistent with the known role of PI3K complex I in autophagy, depletion of Vps34, Vps15, Atg6, or Atg14 led to accumulation of Atg8a/p62-positive aggregates, indicating impaired macroautophagy (Supplementary Fig. 2). In contrast, Uvrag depletion resulted in a milder phenotype, suggesting that defects in lysosomal function also contribute to impaired autophagic degradation.

Together, these results indicate that PI3K complex II, but not the Atg14-containing complex I, is required to maintain the organization of late endosomes and lysosomes in nephrocytes.

### Recruitment of the HOPS tethering complex to late endosomes requires PI3K complex activity

The homotypic fusion and vacuole protein sorting (HOPS) complex is a key tethering complex required for late endosome–lysosome fusion (Nickerson, Brett and Merz 2009, Balderhaar and Ungermann 2013). To test whether recruitment of HOPS is affected by PI3K complex depletion, we examined the localization of the HOPS subunit Vps41 in nephrocytes expressing Vps41-9xHA.

In control and Atg14-depleted nephrocytes, Vps41-9xHA localized to a subset of Rab7-positive endosomes (Supplementary Fig. 3A,B), consistent with recruitment of HOPS to late endosomal compartments. In contrast, in cells depleted of Vps34, Uvrag, Vps15, or Atg6, Vps41-9xHA signal became largely cytosolic and dispersed (Supplementary Fig. 3C-F). Notably, the large Rab7-positive structures present in these cells were largely devoid of Vps41 signal.

These findings indicate that abnormal endosomal compartments forming in the absence of PI3K complex activity fail to recruit the HOPS tethering complex.

### Loss of PI3K complex activity leads to accumulation of abnormal endolysosomal compartments

To determine how PI3K complex depletion affects endolysosomal ultrastructure, we examined nephrocytes by transmission electron microscopy. In control cells, late endosomes appeared as large compartments containing electron-dense material, while lysosomes were typically smaller dense structures (Fig. 2A). In contrast, depletion of Vps34, Uvrag, Vps15, or Atg6 resulted in the accumulation of abnormal endolysosomal compartments (Fig. 2B–E). These structures frequently contained undegraded cytosolic material and displayed complex internal membrane organization, including intraluminal vesicles, tubular membranes, and lamellar structures. Such compartments were markedly enlarged compared to normal endolysosomes and often contained clusters of electron-dense particles resembling ribosomes. These observations indicate that loss of PI3K complex activity leads to the accumulation of aberrant endolysosomal compartments that fail to properly degrade their contents.

**Figure 2.**
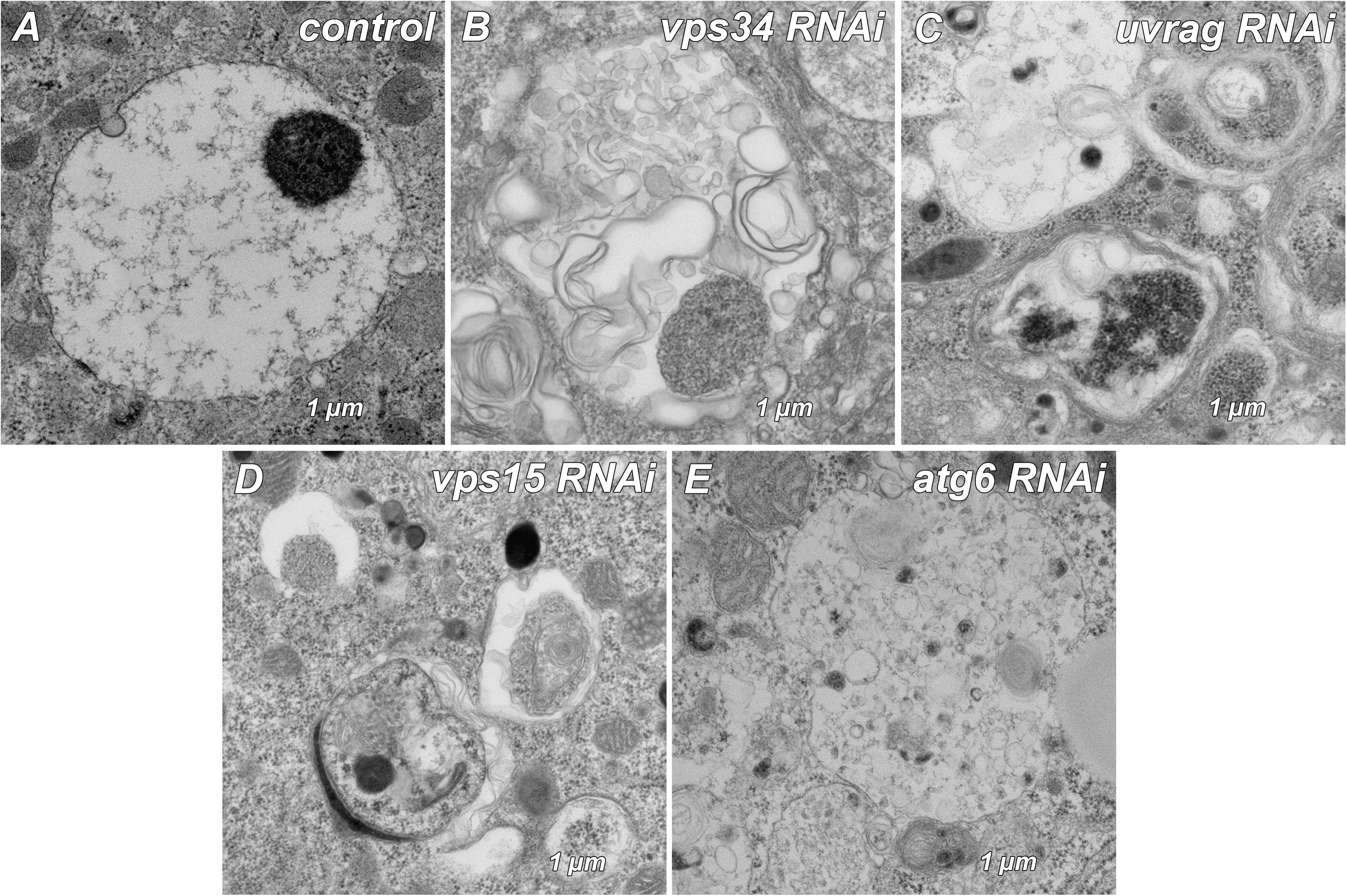
Ultrastructural defects of endolysosomal compartments upon loss of PI3K(III) complex function. Transmission electron microscopy analysis of nephrocytes. **A.** In control cells, late endosomes appear as large compartments containing electron-dense material, while lysosomes are smaller dense structures. **B–E.** In cells depleted of Vps34 (B), Uvrag (C), Vps15 (D), or Atg6 (E), abnormal endosome/lysosome-like compartments accumulate. These structures contain undegraded cytosolic material and multiple intraluminal membrane structures, including vesicles, tubules, and lamellar membranes, indicating defective endolysosomal maturation and degradation.

### Loss of PI3K complex II leads to secretion of abnormal endolysosomal material

Nephrocytes contain a network of plasma membrane invaginations known as lacunar channels, which are connected to the extracellular space and enable rapid uptake and recycling of endocytic cargo (Simons and Huber 2009, Weavers, Prieto-Sanchez et al. 2009, Zhuang, Shao et al. 2009, Helmstädter, Huber and Hermle 2017). Importantly, lacunar channel depth is maintained at a constant level by balancing endosomal degradation and recycling (Wen, Zhang et al. 2020, Hargitai, Molnár et al. 2026). To assess whether PI3K complex II depletion affects this system, we performed channel diffusion assays using fluorescently labeled BSA.

In control nephrocytes, fluorescent BSA labeled the peripheral channel system but did not penetrate deeply into the cells (Fig. 3A). In contrast, nephrocytes depleted of Vps34 or Uvrag displayed significantly increased channel depth (Fig. 3B–C), indicating abnormal expansion of the channel system (Fig. 3D). This is consistent with altered endosomal turnover.

**Figure 3.**
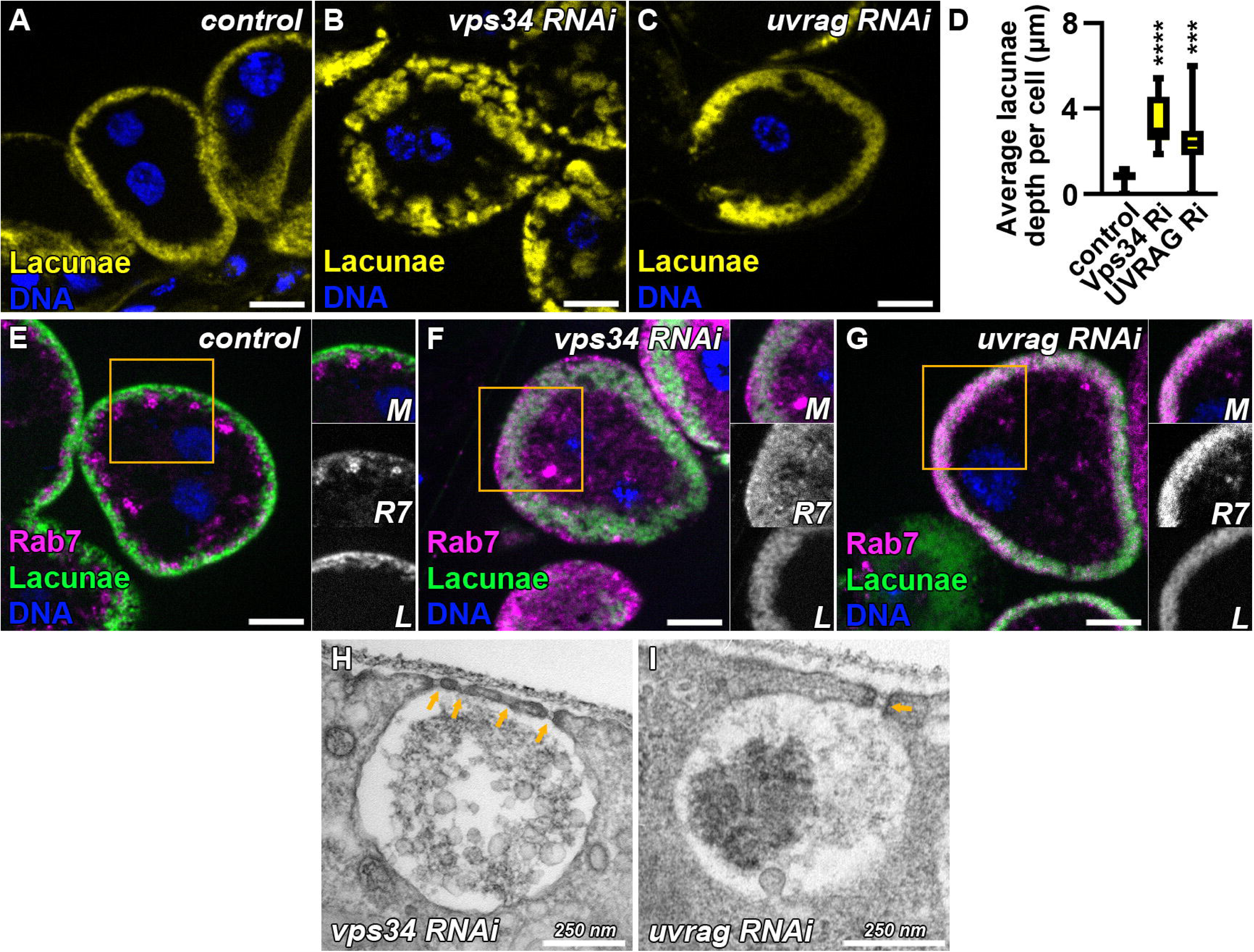
Loss of PI3K(III) complex II promotes lysosome secretion and increases lacunar channel depth. **A–C.** Channel diffusion assay using fluorescently labeled BSA reveals increased lacunar channel depth in nephrocytes expressing Vps34 RNAi (B), Uvrag RNAi (C), compared with control cells (A). Nuclei are shown in blue (DNA). Scale bars: 10 μm. **D.** Quantification of lacunar channel depth shown in panels A–C (n = 15). **E–G.** Following the channel diffusion assay, nephrocytes were stained with anti-Rab7. In Vps34 RNAi (F) and Uvrag RNAi (G) cells, Rab7 signal partially colocalizes with the BSA-labeled lacunar channels compared with controls, suggesting secretion of abnormal late endosomes/lysosomes. Nuclei are shown in blue (DNA). Scale bars: 10 μm. Insets display merged (M), Rab7 (R7), and lacunar channel (L) signals of the boxed regions. **H-I.** Transmission electron microscopy shows endosomal/lysosomal material released into the lacunar channels beneath slit diaphragms (orange arrows) of Vps34 RNAi (H) and Uvrag RNAi (I) nephrocytes.

We also stained nephrocytes for Rab7 after the channel diffusion assay. The Rab7 signal was overlapping with the lacunar channels in Vps34- and Uvrag-depleted cells (Fig. 3E,F,G), suggesting that late endosomal or lysosomal material was released into the channel system. Consistent with this interpretation, transmission electron microscopy revealed endosomal and lysosomal material within the lacunar channels of PI3K complex II–depleted nephrocytes (Fig. 3H-I). These observations indicate that defective endolysosomal maturation is accompanied by secretion of endosomal and lysosomal material in PI3K complex II–deficient cells.

### Inhibition of Hsc70-4 phenocopies the endolysosomal defects of PI3K complex II depletion

The molecular chaperone Hsc70-4 is known to participate in clathrin uncoating and membrane remodeling events during endocytic trafficking (DeLuca-Flaherty, McKay et al. 1990, Cheetham, Anderton and Jackson 1996, Newmyer and Schmid 2001, Chang, Newmyer et al. 2002, Eisenberg and Greene 2007, Böcking, Aguet et al. 2011). Notably, Hsc70-4 mutant nephrocytes have been reported to contain multilamellar lysosome-like structures, resembling those observed upon PI3K(III) depletion (Chang, Newmyer et al. 2002). To test whether Hsc70-4 function might be linked to the PI3K complex II phenotype, we examined nephrocytes expressing a dominant-negative form of Hsc70-4.

Immunostaining for Rab5, Rab7, and Lamp1 revealed that inhibition of Hsc70-4 disrupts endolysosomal organization (Fig. 4A–F). Similar to PI3K complex II depletion, Rab7-positive late endosomes were largely lost, and the remaining Rab7 signal redistributed toward the cell periphery or formed abnormal intracellular structures. Lysosomal organization was similarly affected, closely resembling the phenotype observed upon Vps34 or Uvrag depletion. Quantification confirmed a strong reduction in the size of Rab7-positive endosomes and Lamp1-positive lysosomes (Fig. 4E,F). Transmission electron microscopy further revealed abnormal endolysosomal compartments in nephrocytes expressing dominant-negative Hsc70-4 that were morphologically similar to those observed in PI3K complex-deficient cells (Fig. 4G). To explore whether Hsc70-4 might interact with phosphoinositides, we performed a protein–lipid overlay assay using recombinant Hsc70-4. This analysis revealed that Hsc70-4 binds multiple phosphatidylinositol phosphates, including PI(3)P (Fig. 4H), suggesting that phosphoinositide lipids may contribute to its membrane association.

**Figure 4.**
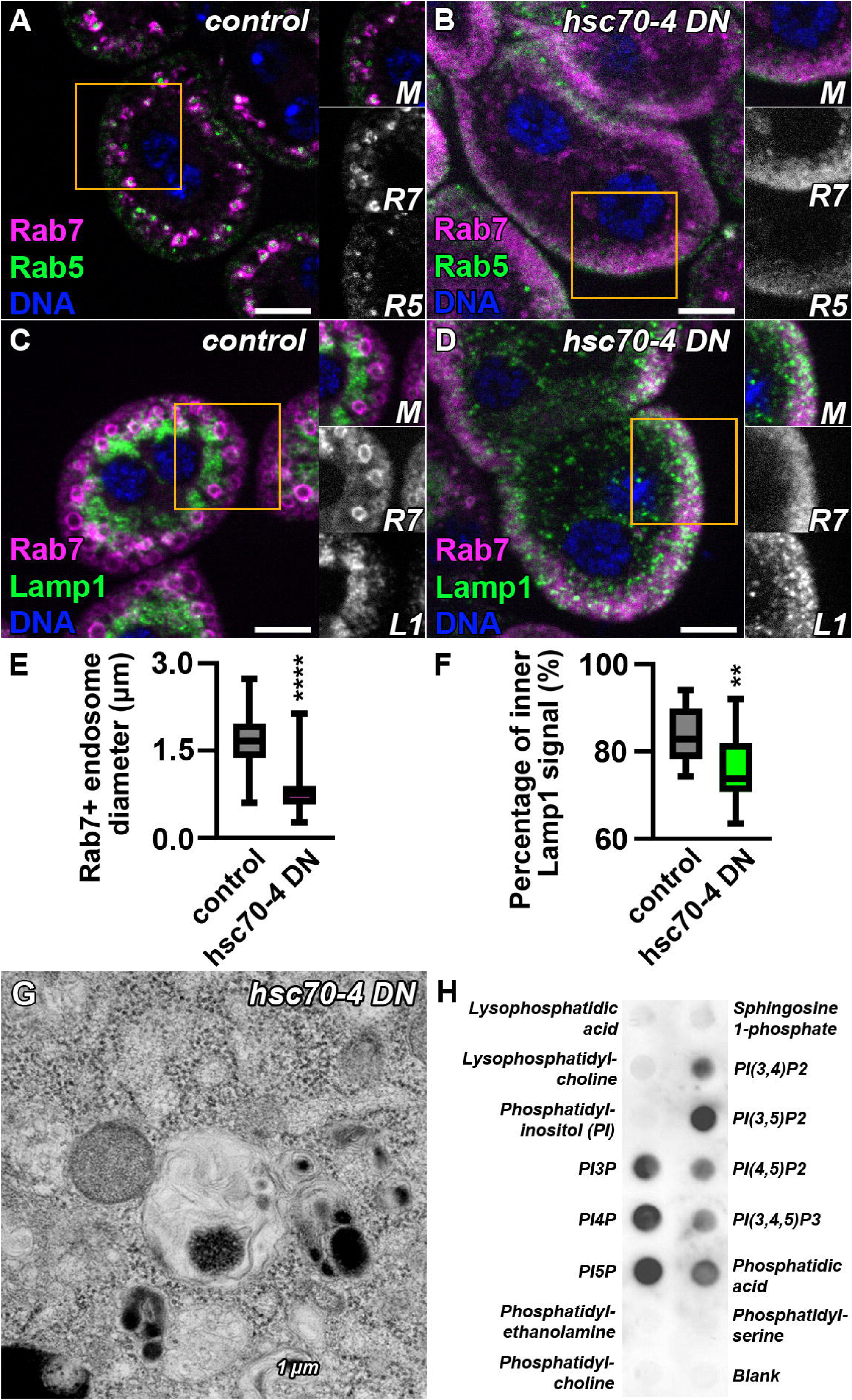
Loss of Hsc70-4 function phenocopies PI3K(III) complex II defects and Hsc70-4 binds phosphoinositides. **A–D.** Nephrocytes were immunostained for Rab7 together with either the early endosomal marker Rab5 (A,B) or the lysosomal marker Lamp1 (C,D). In control cells (A,C), Rab7-positive late endosomes form a distinct layer beneath Rab5-positive early endosomes and above Lamp1-positive lysosomes. In contrast, expression of a dominant-negative Hsc70-4 (Hsc70-4 DN) disrupts this organization (B,D): Rab7-positive late endosomes largely disappear, while Rab7 and Lamp1 signals redistribute toward the cell periphery. In addition, some Rab7-positive structures remain inside the cells. Nuclei are shown in blue (DNA). Scale bars: 10 μm. Insets display merged (M), Rab5 (R5), and Rab7 (R7) or merged (M), Rab7 (R7), and Lamp1 (L1) signals of the boxed regions. **E–F.** Quantification of Rab7-positive endosome size (n = 100) and of the proportion of inner Lamp1 signal (n = 15) shown in panels A–D. **G.** Transmission electron microscopy (TEM) reveals abnormal endosome/lysosome-like structures in Hsc70-4 DN–expressing cells that are similar to those observed in PI3K(III) complex II–depleted cells (Figure 2). **H.** Recombinant Hsc70-4 binds phosphatidylinositol phosphates in a protein–lipid overlay assay using PIP strips. Binding is detected for all phosphatidylinositol phosphates but not for phosphatidylinositol (PI). No binding is observed for lysophosphatidic acid, lysophosphatidylcholine, sphingosine-1-phosphate, phosphatidylethanolamine (PE), phosphatidylserine (PS), or phosphatidylcholine (PC).

Together, these results show that inhibition of Hsc70-4 phenocopies the endolysosomal defects observed upon loss of PI3K complex II.

### Clathrin accumulates on endosomal structures upon loss of PI3K complex II or Hsc70-4

Because Hsc70 proteins are known to mediate clathrin uncoating, we examined whether clathrin distribution is altered in PI3K(III) complex-deficient nephrocytes. To this end, we analyzed nephrocytes expressing GFP-clathrin light chain (GFP-Clc) and stained them for Rab7.

In control cells, GFP-Clc formed a peripheral layer above Rab7-positive late endosomes, consistent with the localization of clathrin-coated endocytic vesicles near the plasma membrane (Fig. 5A). In contrast, in Vps34- or Uvrag-depleted cells, large GFP-Clc–positive structures appeared deeper in the cytoplasm (Fig. 5B,C). Notably, some of these clathrin-positive structures colocalized with Rab7, indicating accumulation of clathrin on abnormal endosomal compartments.

**Figure 5.**
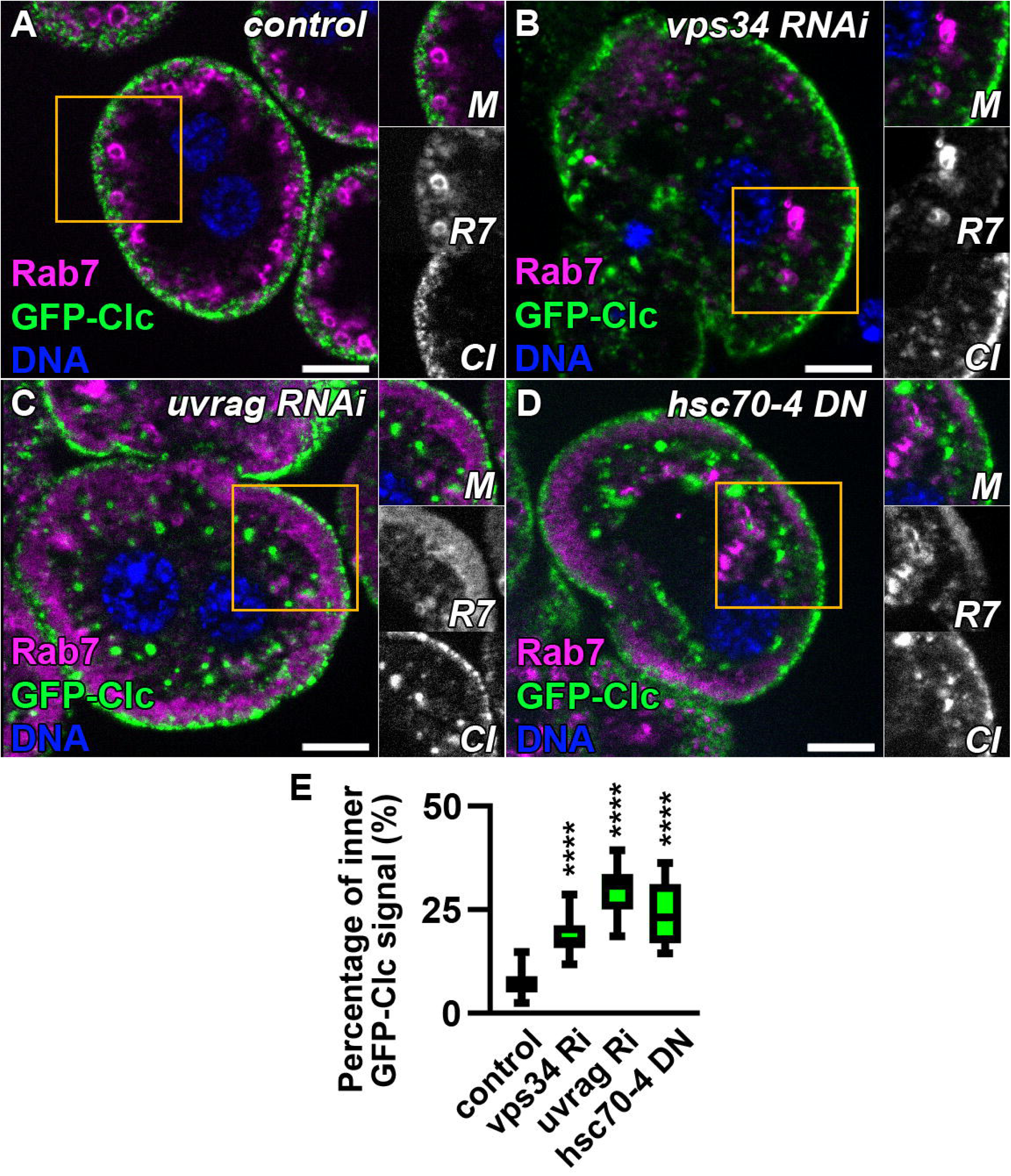
Clathrin accumulates on endosomal structures upon loss of PI3K(III) complex II or Hsc70-4 function. **A–D.** Nephrocytes expressing GFP-clathrin light chain (GFP-Clc) were stained with anti-Rab7 to visualize late endosomes. In control nephrocytes (A), GFP-Clc forms a peripheral layer above Rab7-positive late endosomes. In contrast, in Vps34 RNAi (B) or Uvrag RNAi (C) cells, large GFP-Clc–positive structures appear deeper within the cytoplasm, and some of these structures colocalize with Rab7. A similar phenotype is observed in cells expressing dominant-negative Hsc70-4 (Hsc70-4 DN) (D). Nuclei are shown in blue (DNA). Scale bars: 10 μm. Insets display merged (M), Rab7 (R7), and GFP-Clc (Cl) signals of the boxed regions. **E.** Quantification of inner GFP-Clc signal relative to the total cellular GFP-Clc signal shown in panels A–D.

A similar redistribution of clathrin was observed in cells expressing dominant-negative Hsc70-4 (Fig. 5D). Quantification confirmed a significant reduction of peripheral GFP-Clc signal relative to total cellular clathrin signal in these conditions (Fig. 5E). Additional depletion of other PI3K complex subunits (Vps15 or Atg6) produced similar clathrin redistribution phenotypes, whereas Atg14 knockdown had little effect (Supplementary Fig. 4A–D). Western blot analysis showed comparable GFP-Clc protein levels between control and PI3K complex-depleted larvae, indicating that the observed changes reflect altered clathrin localization rather than altered protein abundance (Supplementary Fig. 4E).

Together, these findings indicate that clathrin abnormally accumulates on endosomal structures when PI3K complex II or Hsc70-4 function is disrupted.

### Clathrin depletion partially rescues endosomal defects caused by loss of PI3K complex II

Because clathrin accumulated on Rab7-positive endosomal structures in PI3K complex II–deficient cells and inhibition of Hsc70-4 produced a similar phenotype, we asked whether persisting clathrin contributes to the endosomal defects observed upon loss of PI3K complex activity. To test this possibility, we depleted clathrin light chain (Clc) alone or in combination with Vps34 knockdown.

Immunostaining for Rab5 and Rab7 revealed that depletion of Clc alone had only minor effects on endosomal organization (Fig. 6A,B). Early and late endosomes maintained their characteristic layered arrangement, although Rab7-positive compartments were slightly enlarged compared with controls. In contrast, nephrocytes expressing both Vps34 RNAi and Clc RNAi displayed a partial restoration of endosomal organization (Fig. 6D–F). Compared with Vps34 single knockdown cells (Fig. 6C), Rab7-positive late endosomes were again detectable within the cytoplasm rather than being restricted to peripheral regions or large structures (Fig. 6D).

**Figure 6.**
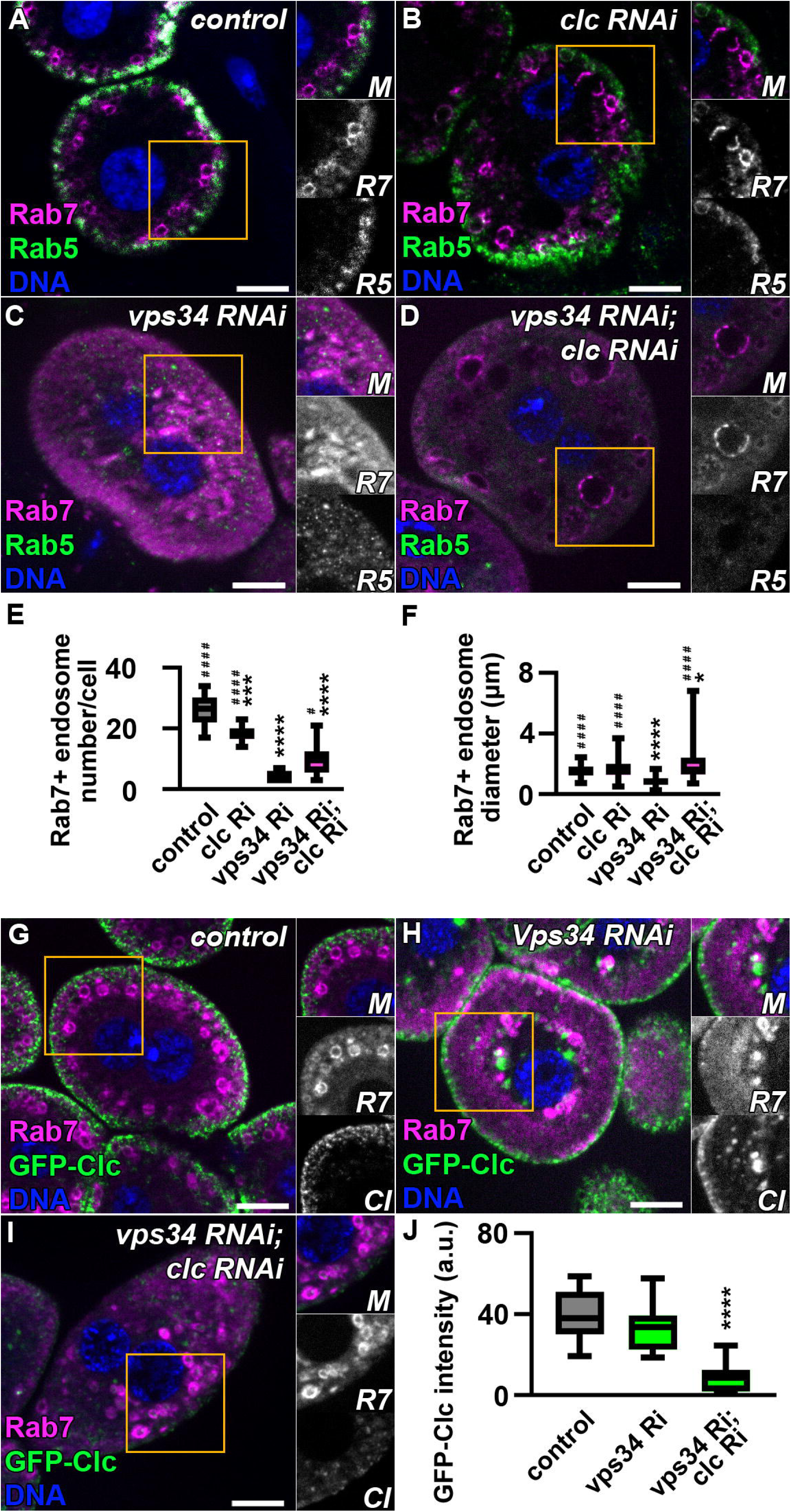
Clathrin depletion partially rescues the endosomal defects caused by Vps34 knockdown. **A–D.** Nephrocytes were immunostained for the early endosomal marker Rab5 and the late endosomal marker Rab7. In control cells (A), Rab5-positive early endosomes form a peripheral layer, while Rab7-positive late endosomes are located in a distinct layer deeper in the cytoplasm. Depletion of clathrin light chain (Clc RNAi) resulted in a slight increase in the size of late endosomes but otherwise did not markedly affect this organization (B). In contrast, in nephrocytes expressing both Vps34 RNAi and Clc RNAi (D), endosomal organization was partially restored compared with Vps34 single knockdown cells (C). Rab7-positive late endosomes reappeared within the cytoplasm, although their size remained significantly larger than in control cells. Nuclei are shown in blue (DNA). Scale bars: 10 μm. Insets display merged (M), Rab5 (R5) and Rab7 (R7) signals of the boxed regions. **E-F.** Quantification of Rab7-positive endosome number (E) and size (F) shown in panels A–D (n = 10). **G–I.** Nephrocytes expressing GFP-Clc were immunostained for Rab7. In control cells (G), GFP-Clc localizes to peripheral clathrin-coated endocytic vesicles. Expression of Vps34 RNAi did not eliminate the GFP-Clc signal but did alter Rab7-positive late endosomes (H). In Vps34 and Clc double RNAi cells (I), clathrin signal is absent, and Rab7-positive late endosomes are present within the cytoplasm, indicating partial restoration of late endosomal organization. Nuclei are shown in blue (DNA). Scale bars: 10 μm. Insets display merged (M), Rab7 (R7), and GFP-Clc (Cl) signals of the boxed regions. **J.** Quantification of GFP-Clc signal shown in panels G–I (n = 15).

To confirm that this rescue was due to clathrin depletion, we analyzed nephrocytes expressing GFP-Clc (Fig. 6G–J). Expression of Clc RNAi efficiently eliminated the GFP-Clc signal in Vps34 RNAi backgrounds (Fig. 6I,J), confirming robust depletion of clathrin. In Vps34 and Clc double knockdown cells, Rab7-positive late endosomes were present within the cytoplasm, and the large clathrin- and Rab7-positive structures observed in Vps34 single knockdown cells (Fig. 6H) were largely absent (Fig. 6I).

Together, these results indicate that persisting clathrin contributes to the endosomal defects caused by PI3K complex II depletion and that reducing clathrin levels partially alleviates these defects.

Consistent with a functional connection between PI3K complex II and clathrin removal, Hsc70-4, a chaperone involved in clathrin uncoating, showed partial spatial overlap with clathrin-positive structures in nephrocytes. Although the majority of Hsc70-4 signal was cytosolic, enrichment was detectable near the cell periphery where clathrin-coated endocytic vesicles are located (Supplementary Fig. 5A,B). Upon depletion of Vps34, this peripheral enrichment was reduced and Hsc70-4 signal appeared more uniformly distributed throughout the cytosol (Supplementary Fig. 5C), suggesting that PI3K(III) complex activity contributes to the spatial organization of Hsc70-4 in nephrocytes.

Taken together, these findings support a model in which the Uvrag-containing PI3K complex promotes endosomal maturation at least in part by enabling Hsc70-4–dependent removal of clathrin from endosomal membranes.

## Discussion

Phosphatidylinositol 3-kinase III (PI3K(III)) plays a central role in endolysosomal trafficking by generating phosphatidylinositol-3-phosphate (PI(3)P), a lipid that contributes to endosomal and autophagosomal membrane identity and recruits numerous effector proteins (Kihara, Noda et al. 2001, Backer 2008, Itakura, Kishi et al. 2008, Funderburk, Wang and Yue 2010). While the Uvrag-containing PI3K complex II has long been implicated in endosome maturation and lysosomal function, the mechanistic basis of this requirement has remained incompletely understood. Here, using Drosophila garland nephrocytes, we identify a previously unrecognized role for PI3K complex II in promoting the removal of clathrin from endosomal membranes. Loss of complex II results in persistent clathrin association with Rab7-positive compartments, disruption of late endosomal organization, defective recruitment of the HOPS tethering complex, and impaired lysosomal maturation. These defects are closely phenocopied by inhibition of the clathrin uncoating chaperone Hsc70-4, and partial depletion of clathrin alleviates the Vps34 knockdown phenotype. Together, these findings indicate that PI3K complex II supports endolysosomal maturation at least in part by enabling Hsc70-4–dependent clathrin removal.

Previous studies have demonstrated that PI3K complex II components such as Uvrag are required for endosomal maturation and lysosomal degradation, but how PI(3)P production contributes to these processes has remained unclear (Juhász, Hill et al. 2008, Abe, Setoguchi et al. 2009, Lee, Liang et al. 2011, Lorincz, Lakatos et al. 2014, Takats, Pircs et al. 2014, Munson, Allen et al. 2015). In PI3K complex II–deficient nephrocytes, we observed a collapse of endolysosomal organization, including altered Rab7 compartments, disorganized Lamp1-positive lysosomes, and accumulation of abnormal endolysosomal structures. These compartments failed to recruit the HOPS tethering complex, consistent with defective acquisition of late endosomal identity. Our findings therefore provide a mechanistic framework linking PI3K complex II activity to the establishment of functional late endosomes capable of undergoing normal fusion and maturation.

A central observation of this study is that clathrin abnormally accumulates on endosomal structures in the absence of PI3K complex II activity. While clathrin is best known for its role in endocytic vesicle formation, it also functions on endosomal membranes and must be efficiently removed to allow proper membrane remodeling. Clathrin association and removal are tightly regulated during lysosome reformation, where clathrin-dependent tubulation and scission are required to regenerate functional lysosomes. Both insufficient and persistent clathrin association can impair this process and lead to the accumulation of dysfunctional lysosomes (Rong, Liu et al. 2012, McGrath, Eramo et al. 2021, Nanayakkara, Gurung et al. 2023). In addition, phosphoinositides, including PI(3)P, have been implicated in lysosome reformation (Munson, Allen et al. 2015, Khundadze, Ribaudo et al. 2021). Our data suggest that loss of PI3K(III) complex II disrupts this balance by preventing efficient clathrin removal, thereby impairing endosomal maturation and potentially lysosome reformation.

Consistent with this model, inhibition of Hsc70-4, a key mediator of clathrin uncoating (DeLuca-Flaherty, McKay et al. 1990, Cheetham, Anderton and Jackson 1996, Newmyer and Schmid 2001, Chang, Newmyer et al. 2002, Eisenberg and Greene 2007, Böcking, Aguet et al. 2011), phenocopies PI3K complex II depletion, and clathrin accumulates on intracellular - often Rab7-positive - compartments in both conditions. Moreover, partial depletion of clathrin alleviates the defects caused by Vps34 knockdown, indicating that persistent clathrin is functionally relevant rather than a secondary consequence. We also observed that Hsc70-4 shows partial enrichment at clathrin-positive structures and that this spatial organization is altered upon Vps34 depletion. Together with the ability of Hsc70-4 to bind phosphoinositides, these findings suggest that PI(3)P-enriched membranes may facilitate Hsc70-4–dependent clathrin removal during endosomal maturation.

Lysosomal exocytosis provides an alternative route for cells to eliminate undegraded material when lysosomal function is compromised. In this process, lysosomes fuse with the plasma membrane and release their contents into the extracellular space (Medina, Fraldi et al. 2011, Samie and Xu 2014, Buratta, Tancini et al. 2020, Tancini, Buratta et al. 2020, Domingues, Catarino et al. 2024). Consistent with this, we observed that PI3K complex II–deficient nephrocytes release endolysosomal material into lacunar channels, suggesting that lysosomal exocytosis may act as a compensatory mechanism in response to impaired degradation.

An additional intriguing feature of the abnormal endolysosomal compartments observed in PI3K complex-deficient nephrocytes is the presence of cytosolic material within their lumen. Electron microscopy frequently revealed ribosome-rich cytosolic content within these compartments, whereas larger organelles such as mitochondria or endoplasmic reticulum membranes were rarely observed. Although these structures resemble amphisome-like intermediates, the selective accumulation of small cytosolic material suggests that cargo uptake may occur through a mechanism distinct from canonical macroautophagy. One possibility is endosomal microautophagy, a process in which endosomal membranes directly internalize cytosolic components and which involves Hsc70 family chaperones (Sahu, Kaushik et al. 2011, Uytterhoeven, Lauwers et al. 2015, Tekirdag and Cuervo 2018, Mesquita, Glenn and Jenny 2020). Whether endosomal microautophagy contributes to the formation of the cytosol-containing compartments observed here remains an interesting question for future studies.

In summary, our findings reveal a functional link between PI3K complex II activity, clathrin dynamics, and late endosomal maturation. We propose that PI3K complex II–generated PI(3)P enables Hsc70-4–dependent removal of clathrin from endosomal membranes, thereby allowing recruitment of the HOPS tethering complex and progression toward functional lysosomes. When this pathway is disrupted, clathrin persists on endosomal membranes, vesicle maturation is impaired, and abnormal endolysosomal compartments accumulate that can be secreted. These results uncover a new role for PI3K complex II in regulating membrane remodeling during endolysosomal trafficking.

## Materials and Methods

### Fly work

The flies were kept in glass tubes on a standard medium which contained cornmeal, yeast and sucrose, and were raised at 25°C. The following RNA interference stocks for Vps34 (33384) (FlyBase ID: FBst0033384), Uvrag (34368) (FlyBase ID: FBst0034368), Vps15 (34092) (FlyBase ID: FBst0034092), Atg6 (28060) (FlyBase ID: FBst0028060), Atg6 (35741) (FlyBase ID: FBst0035741), white[1118] (3605) (FlyBase ID: FBst0003605), luciferase (31603) (FlyBase ID: FBst0031603),Hsc70-4 dominant-negative mutant (5845) (FlyBase ID: FBst0005845), UAS-Clc-GFP (7109) (FlyBase ID: FBst0007109), UAS-GFP-myc-2xFYVE (42712) (FlyBase ID: FBst0042712) and prospero-Gal4 driver (80572) (FlyBase ID: FBst0080572) were obtained from the Bloomington Drosophila Stock Center (BDSC), Bloomington, IN, USA. The RNAi lines for Vps34 (v100296) (FlyBase ID: FBst0472170), Atg6 (v110197) (FlyBase ID: FBst0481778), Atg6 (v22123) (FlyBase ID: FBst0454427) and Atg14 (v108559) (FlyBase ID: FBst0480369) were obtained from Vienna Drosophila Resource Center (VDRC), Vienna, Austria. The UAS-Vps41-9xHA riporter was already in our hands (Lorincz, Kenez et al. 2019). The UAS-HA-Hsc70-4^WT^ was a kind gift from Ioannis P. Nezis (Jacomin, Gohel et al. 2021).

All genotypes used in the experiments can be found in Table 1.

### Immunohistochemistry

For immunostainings, garland nephrocytes of stage L3 wandering larvae were dissected in PBS (Sigma-Aldrich, cat#: P38135) and fixed for 40 minutes in 4% formaldehyde (Sigma-Aldrich, cat#: 252549) in PBS at room temperature. Next, the samples were incubated in PBS for 10 minutes, then permeabilized in 0,1 % PBTX (PBS containing 0,1% Triton X-100 (Sigma-Aldrich, cat#: X100)). Then garland cells were blocked in PBTX containing 5% fetal bovine serum (Sigma-Aldrich, cat#: F4135) for 30 minutes. After that samples were transferred into blocking solution containing the primary antibodies and incubated overnight at 4°C. We next washed the cells 3 times with PBTX and incubated them in PBTX containing 4% NaCl (Sigma-Aldrich, cat#: S5886) for 10 minutes, after that samples were rinsed in PBTX and incubated in PBTX for 2×15 minutes. Nephrocytes were incubated for 30 minutes in blocking solution and in secondary antibodies dissolved in blocking solution for 3 hours. Then the samples were rinsed 3 times in PBTX and washed for 15 minutes in PBTX containing 4% NaCl and 5µg/ml Hoechst 33342 staining (Thermo Fisher Scientific, cat#: 62249). After that the cells were rinsed in PBTX, incubated 2×15 minutes in PBTX, washed 3 times in PBS and incubated 2×15 minutes in PBS. Finally, samples were mounted in Vectashield Antifade Mounting Medium (Vector Laboratories, H-1000-10).

The following antibodies were used: monoclonal mouse anti-Rab7 (1:10, DSHB, RRID: AB_2722471), polyclonal rabbit anti-Rab5 (1:100, Abcam, RRID: AB_882240), polyclonal rabbit anti-HA (1:100, Merck, RRID: AB_260070), polyclonal rat anti-Atg8a (1:300, (Takats, Nagy et al. 2013)), polyclonal rabbit anti-p62 (1:1000,(Pircs, Nagy et al. 2012)), polyclonal rabbit anti-Lamp1 (1:1000,(Chaudhry, Sica et al. 2022)) and polyclonal guinea pig anti-GFP (1:500, (Takats, Nagy et al. 2013)), Alexa Fluor^TM^ 488 goat anti-rabbit (1:1000, Thermo Fisher Scientific, RRID: AB_2630356), Alexa Fluor^TM^ 568 donkey anti-mouse (1:1000, Thermo Fisher Scientific, RRID: AB_11180865), Alexa Fluor^TM^ 568 goat anti-rat (1:1000, Thermo Fisher Scientific, RRID: AB_2534121) and DyLight 488 anti-guinea pig (1:600, Invitrogen, cat#: SA5-10094).

### Electron microscopy

For electron microscopy, dissected garland nephrocytes were fixed in 3.2% paraformaldehyde (Sigma-Aldrich, cat#: P6148), 1% glutaraldehyde (Sigma-Aldrich, cat#: G7776), 1% sucrose (Sigma-Aldrich, cat#: S0389) and 0.003 M CaCl_2_ (Sigma-Aldrich, cat#: C1016) containing 0.1 M sodium cacodylate buffer (pH 7.4, Sigma-Aldrich, cat#: C0250) overnight at 4°C. Next the samples were postfixed in 0.5% osmium tetroxide (Sigma-Aldrich, cat#: O5500) for 1 hour and incubated in half-saturated aqueous uranyl acetate (TAAB, ref#: U001/S/2/10) for 30 minutes. After that, the specimens were dehydrated in a graded series of ethanol (Novochem, cat#: 3C84L0HL120) and embedded into Durcupan ACM (Sigma-Aldrich, cat#: 44614), according to the manufacturer’s recommendations. 70 nm sections of the samples were stained with Reynold’s lead citrate (Thermo Fisher Scientific, cat#: A10701.22) (Lorincz, Lakatos et al. 2016).

### Channel diffusion assay

For the channel diffusion assay garland nephrocytes were fixed in 4% formaldehyde (Sigma-Aldrich, cat#: 252549) in PBS (Sigma-Aldrich, cat#: P38135) for 5 minutes and incubated for 15 minutes (room temperature) in Alexa Fluor 647-bovine serum albumin (Thermo Fisher Scientific, cat#: A34785) dissolved in Shield and Sang M3 Insect Medium (Sigma-Aldrich, cat#: S8398). Next, the samples were postfixed for 45 minutes in 4% formaldehyde in PBS and washed in PBS for 3×10 minutes (Milosavljevic, Lempicki et al. 2022). Then the samples were mounted in Vectashield Antifade Mounting Medium (Vector Laboratories, H-1000-10).

### Molecular Cloning

The following primers were used to establish N-terminally 6xHis-tagged Hsc70-4: Forward: 5′-ACAGCCAGGATCCGAATTCGGGCGGAGGTGGTTCTATG TCTAAAGCTCCTGCTGTTGG-3′; Reverse: 5′-GCGGCCGCAAGCTTGTCGACTTAGTCGACCTCCTCGATGGTGG -3′. Clone LP19893 was used as a template (Berkeley Drosophila Genome Project; LP CDNA library) to amplify full length Hsc70-4 sequence, and the PCR product was inserted to EcoRI-SalI site of pET Duet vector using NEBuilder HiFi DNA Assembly Cloning Kit.

### Recombinant protein expression and purification

6xHis-Hsc70-4 protein were produced in BL21(DE3) bacterial strain following induction with 0,1 mM isopropyl β–D-thiogalactopyranoside (IPTG) for 16 h at 18°C. Cells were harvested by centrifugation, resuspended in buffer containing 20 mM Tris, 150 mM NaCl, 0,1% Triton-X 100, 2 mM beta-mercaptoethanol, pH=7,5 and lysed by sonication on ice. The soluble fraction was obtained by centrifugation at 20 000 rpm for 30 min at 4°C. The fusion proteins were immobilized on Ni Sepharose™ excel (Cytiva) beads for 2 hours at 4°C, washed with the following buffers: 50 mM Na_2_HPO_4_, 600 mM NaCl, pH=7,5 and 50 mM Na_2_HPO_4,_ 300 mM NaCl, 40 mM imidazol, pH=8,0 and eluted with 50 mM Na_2_HPO_4,_ 300 mM NaCl, 400 mM imidazol, pH=8,0. To increase protein concentration and for buffer exchange, Amicon® Ultra-15 (30 000 MWCO, Millipore) concentration tube was used and the protein was stored in 20 mM Tris, 150 mM NaCl, 1 mM ditiotreitol, 10% glycerol, pH=7.5.

### Lipid strip assay

Lipid strip assay was performed according to the manufacturer instructions (PIP Strips™ membrane, Invitrogene). Briefly, PIP Strips membranes were blocked with 10 mM Tris–HCl, pH = 8,0, 150 mM NaCl, 0,1% Tween20, 3% fatty acid free BSA and then incubated with 0,5 μg/ml 6xHis-Hsc70-4 protein (diluted in blocking buffer) for 16 hours at 4°C. To detect bound proteins, anti-polyHistidine tag antibody (Merck, H1029) diluted in 1:5000 and horseradish peroxidase (HRP)-conjugated anti-mouse antibody (Merck, A9044) 1:8000 was used, followed by incubation with Immobilon Western Chemiluminescent HRP Substrate (Millipore).

### Western blot

Western blot experiments were performed as in the previously described protocol (Takats, Nagy et al. 2013). L3 wandering larvae were collected in 1:1 PBS (Sigma-Aldrich, cat#: P38135) and Laemmli-buffer mixture and boiled for 5 minutes, then after homogenization, the samples were again boiled for 5 minutes. Next, the samples were centrifuged for 5 minutes at 13300 rpm. Then the middle protein-rich fraction was used for western blot analysis.

For protein separation 12% acrylamide gels containing SDS (sodium dodecyl sulfate, Sigma-Aldrich, cat# 151-21-3) were used. The running buffer consisted of 0.025M Trizma base (Sigma-Aldrich, cat# T1503), 0.05% SDS and 0.2M glycine (VWR International, cat# 24403.298) dissolved in distilled water. Trans-Blot Turbo Transfer System (Bio-Rad Laboratories) was used to transfer protein samples to Immun-Blot PVDF membranes (Bio-Rad Laboratories, cat#1620174). The buffer used for the transfer consisted of 20% methanol, 0.025M Trizma base, 0.05% SDS and 0.192M glycine dissolved in distilled water. Then membranes were blocked overnight at 4 °C in 0,5% casein (Sigma-Aldrich, cat# C3400) dissolved in TBS (Tris-Buffered Saline). After that, membranes were incubated in primary antibodies for 1 hour in 1:1 -casein TBST mixture. TBST consisted of TBS and 0.1% Tween20 (Sigma-Aldrich, P7949). Next, membranes were washed three times for ten minutes in TBST. Then, membranes were incubated for one hour with secondary antibodies dissolved in casein-TBST 1:1 mixture and again washed three times for ten minutes in TBST. To detect proteins, the DAB peroxidase substrate kit (Vector Laboratories, cat# SK-4100) was used.

These primary antibodies were used: polyclonal guinea pig anti-GFP (1:3000, (Takats, Nagy et al. 2013), and monoclonal mouse anti-tubulin (1:2000, DSHB, cat# AA4.3). The following secondary antibodies were used: horseradish peroxidase (HRP)-conjugated anti-mouse antibody (1:10000, Merck, A9044) and horseradish peroxidase (HRP)-conjugated anti-guinea pig antibody (1:2500).

### Imaging

Fluorescent images were taken with an AxioImager M2 microscope (Carl Zeiss), equipped with an Apotome2 grid confocal unit (Carl Zeiss). We used a Plan-Apochromat 63×/1.40 Oil objective (Carl Zeiss), and an Orca Flash 4.0 LT sCMOS camera (Hamamatsu Photonics) Images were processed with the ZEN 2.3 software (Carl Zeiss). Electron microscopic images were taken with a JEOL JEM-1011 transmission electron microscope equipped with a Morada camera (Olympus) and iTEM software (Olympus). Photoshop CS6 (Adobe) were used for creating figures.

### Quantification and Statistics

ImageJ software (National Institutes of Health) was used for quantification of fluorescent structures or structures were quantified manually. The signal threshold was set by the same person per experiment with ImageJ. For quantification original, single sections were used and cells with nuclei in the focal plane were chosen randomly.

For GFP-FYVE signal intensity measurements (Fig. 1Q) pictures were chosen from 3-4 animals, and at least 4-5 cells were selected from each animal and 44 GFP-FYVE positive structures’ integrated density were measured. Then areas next to the GFP-FYVE positive structures were selected, and their integrated density were measured. After that, the ratio of the two integrated density values was calculated.

ImageJ line tool was used for measuring Rab7+ endosome diameter (Fig. 1R, Fig. 4E, Fig. 6F, Supplementary Fig. 1N) and TGN-YFP width (Fig. 1T). For each genotype 3-5 animals, 10-10 cells and 100-100 fluorescent structures were quantified.

For quantification of GFP-Clc signal intensity (Fig. 6J) each cell’s area and integrated density were measured with ImageJ. Then the integrated density/area ratio was calculated. For each genotype pictures of 4-6 animals were used and 15 cells were quantified.

Atg8a or p62 area fractions of cells were measured with ImageJ for Atg8a and p62 coverage quantification (Supplementary Fig. 2G,H). 3 to 5 animals were used per genotype, and a total of 15-15 cells were analyzed.

For quantification of the percentage of inner Lamp1 (Fig. 1S, Fig. 4F) or GFP-Clc (Fig. 5E) signal whole cell’s and enlarged section’s integrated density was measured with ImageJ. In case of Lamp1 the enlarged section was 1,5 μm smaller, in case of GFP-Clc it was 5 μm smaller. Then integrated density ratio of inner section/whole cell was calculated. For each genotype 5 animals were used and 15 cells were counted.

To analyze lacunar depth (Fig. 3D), the total cell area and lacunar endpoint area were determined. Then the difference of the radius of the calculated circles was used to quantify an approximate lacunar depth. For measurements 15 cells of 3-4 animals were used.

The number of Rab7 positive endosomes was quantified manually. For each genotype 3-5 animals were used, and 10-10 cells were counted (Fig. 6E)

For statistical analyses GraphPad Prism 9.4.1 (GraphPad) was used. Normality was determined with the D’Agostino-Pearson test. Table 2. contains all used tests and post hoc tests for each statistics panel.

## ACKNOWLEDGMENTS

We thank S. Pálfia, M. Truszka, and I. Répássy for technical assistance and colleagues listed in Materials and Methods for providing reagents.

## Data and materials availability

All data needed to evaluate the conclusions in the paper are present in the paper and/or the Supplemental Materials.

## Author contributions

**A.N.:** Conceptualization, Formal analysis, Funding acquisition, Investigation, Methodology, Validation, Visualization and Writing – original draft, Writing – review & editing. **V.B.:** Formal analysis and Investigation. **D.H.:** Formal analysis and Investigation. **A.B.:** Formal analysis and Investigation. **E.H.:** Formal analysis and Investigation. **Z.S-V.:** Formal analysis, Methodology and Investigation. **G.J.:** Funding acquisition, Methodology and Resources, Writing – review & editing. **P.L.:** Conceptualization, Funding acquisition, Investigation, Methodology, Project administration, Resources, Supervision, Validation, Visualization and Writing – original draft, Writing – review & editing.

## Figure legends

**Supplementary Figure 1.**
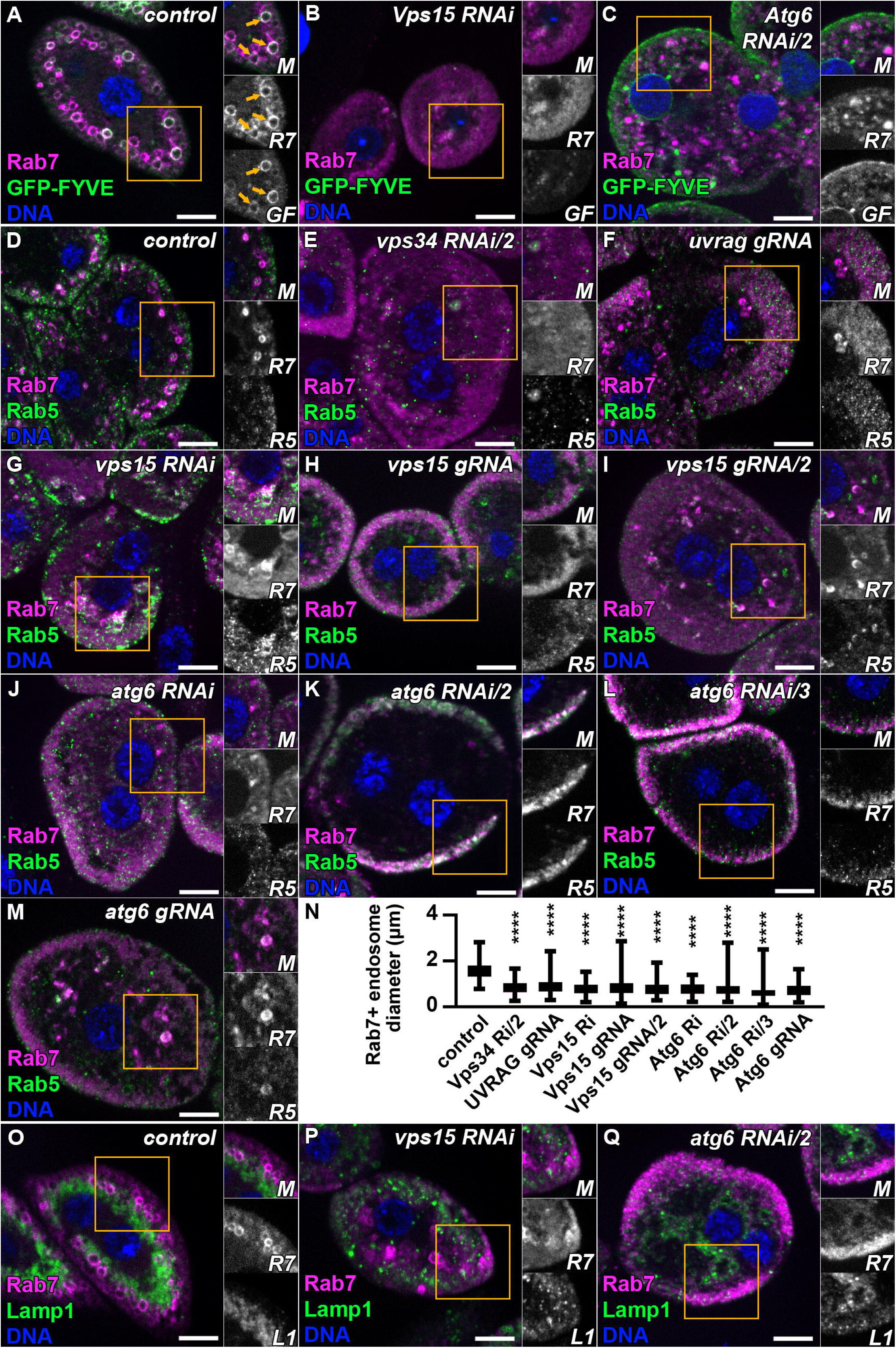
Additional PI3K(III) complex components are required for normal endosomal organization. **A–C.** Nephrocytes expressing GFP-FYVE (A) and RNAi constructs targeting PI3K(III) regulatory subunits Vps15 (B) or Atg6 (C), immunostained for Rab7, display phenotypes similar to those observed upon depletion of Vps34 or Uvrag in Figure 1, including loss of PI(3)P signal and abnormal Rab7-positive compartments. Nuclei are shown in blue (DNA). Scale bars: 10 μm. Insets display merged (M), Rab7 (R7), and GFP-FYVE (GF) channels. **D–M.** Nephrocytes expressing independent RNAi constructs or gRNAs targeting Vps34 (E), Uvrag (F), or the PI3K(III) regulatory subunits Vps15 (G–I) and Atg6 (J–M) were immunostained for Rab7 together with the early endosomal marker Rab5. All manipulations produced phenotypes comparable to those observed for Vps34 or Uvrag depletion in Figure 1. Nuclei are shown in blue (DNA). Scale bars: 10 μm. Insets display merged (M), Rab5 (R5) and Rab7 (R7) signals of the boxed regions. **N.** Quantification of Rab7-positive endosome size shown in panels D–M (n = 100). **O–Q.** Nephrocytes expressing RNAi constructs targeting PI3K(III) regulatory subunits Vps15 (P) and Atg6 (Q) were immunostained for Rab7 together with Lamp1. These cells show phenotypes comparable to those observed for Vps34 or Uvrag depletion in Figure 1. Nuclei are shown in blue (DNA). Scale bars: 10 μm. Insets display merged (M), Rab7 (R7), and Lamp1 (L1) signals of the boxed regions.

**Supplementary Figure 2.**
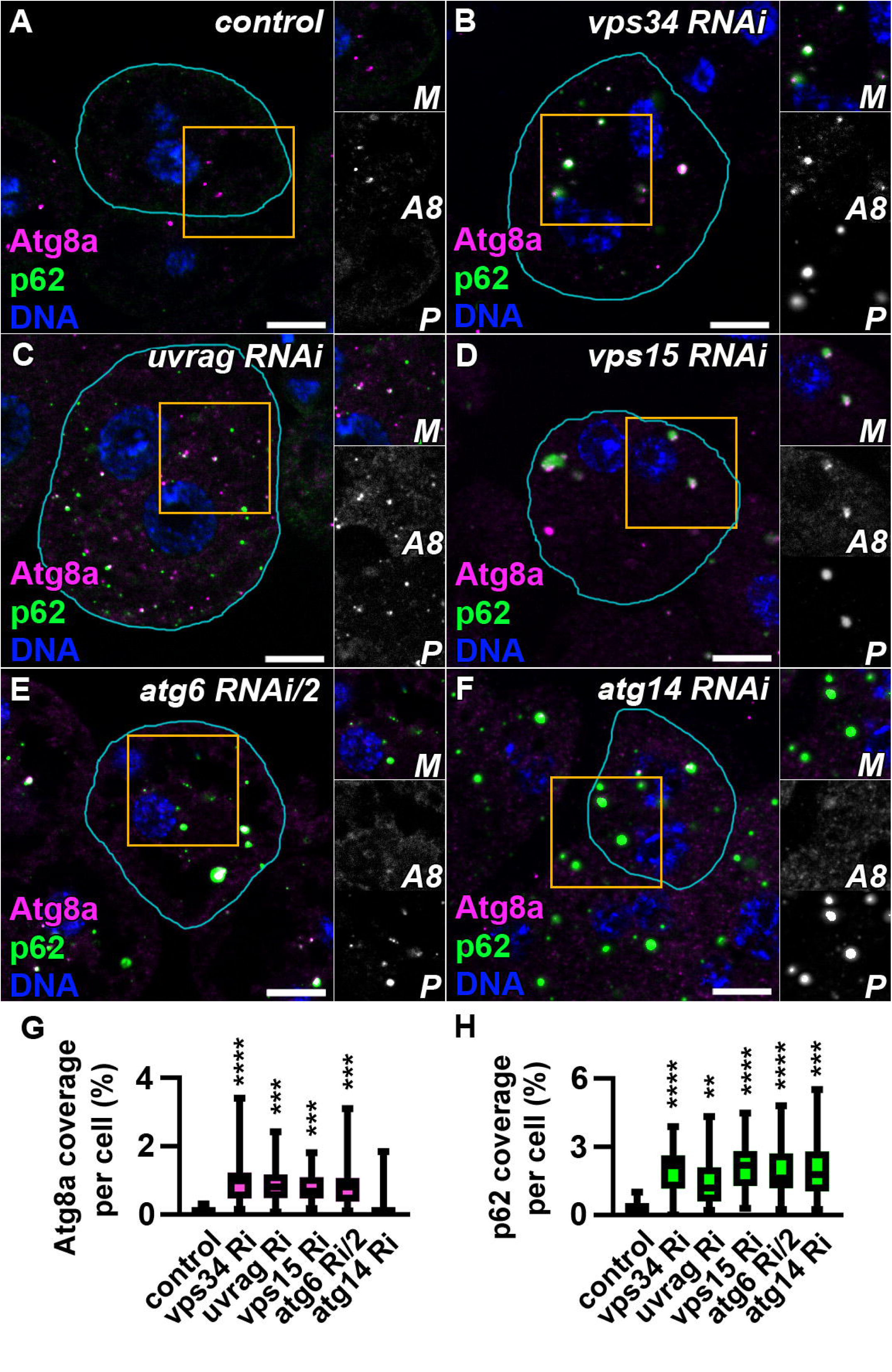
PI3K(III) complex depletion disrupts autophagy in nephrocytes. **A–F.** Nephrocytes were stained for Atg8a and Ref2P/p62. Compared with control cells (A), large Atg8a/p62 double-positive aggregates accumulate in Vps34 RNAi (B), Vps15 RNAi (D), Atg6 RNAi (E), and Atg14 RNAi (F) cells, indicating defective macroautophagy. Uvrag-depleted cells (C) also display increased Atg8a/p62-positive structures, although to a lesser extent. Nuclei are shown in blue (DNA). Scale bars: 10 μm. Insets display merged (M), Atg8a (A8) and p62 (P) signals of the boxed regions. **G–H.** Quantification of Atg8a/p62 coverage shown in panels A–F. Uvrag-deficient cells contain significantly more p62 aggregates than controls but fewer than other PI3K(III) component knockdowns, suggesting that impaired lysosomal function contributes to reduced autophagic degradation, albeit less strongly than defects affecting the autophagosome-generating complex (n = 15).

**Supplementary Figure 3.**
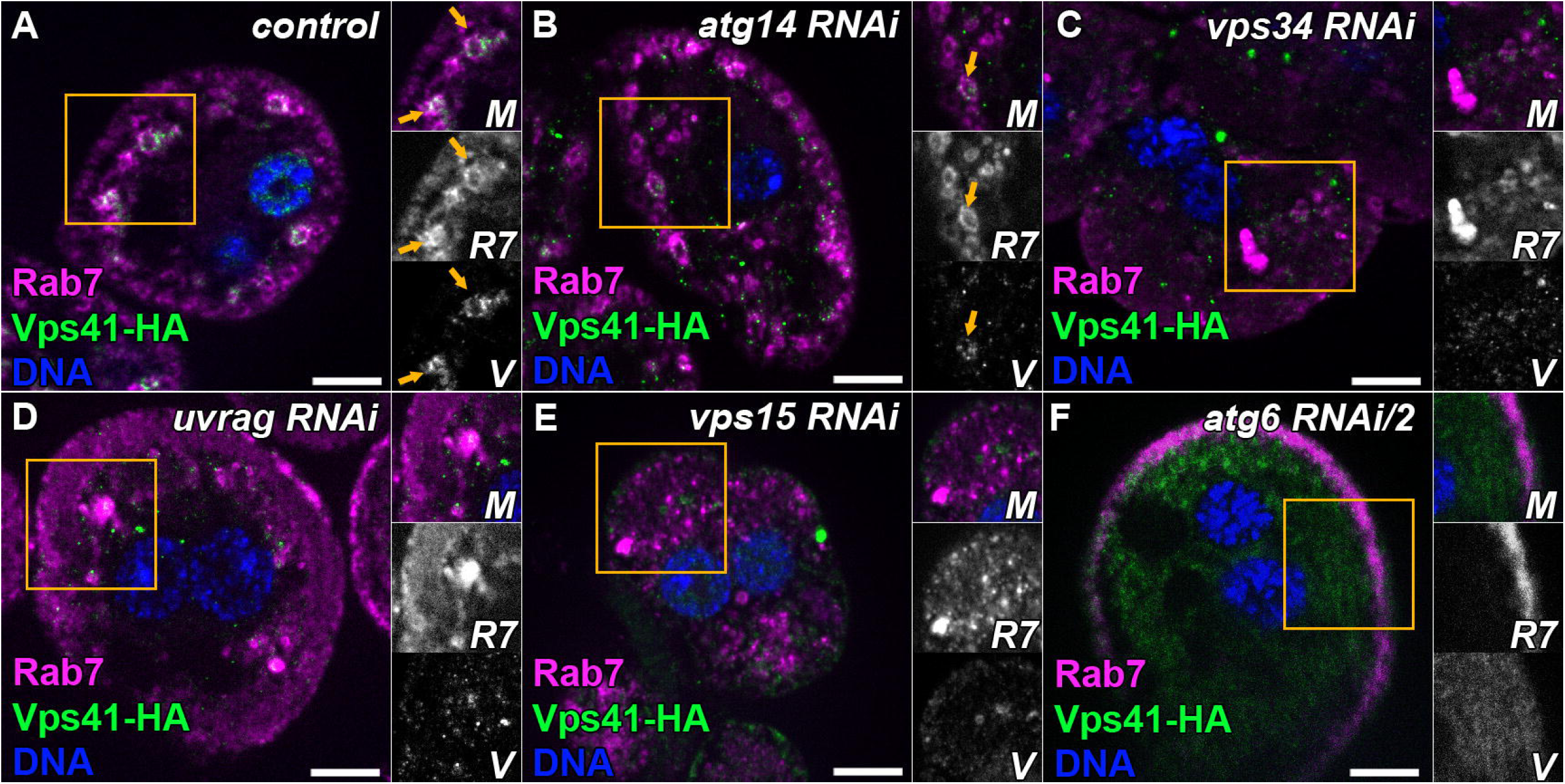
Recruitment of the HOPS subunit Vps41 to late endosomes requires PI3K(III) complex function. **A-F.** All images show nephrocytes expressing Vps41-9xHA in different genetic backgrounds and stained with anti-HA (green) and anti-Rab7 (magenta). Nuclei are shown in blue (DNA). Scale bars: 10 μm. Insets display merged (M), Rab7 (R7), and Vps41-9xHA (V) signals of the boxed regions. **A–B.** In control and Atg14 RNAi nephrocytes, Vps41-9xHA is recruited to a subset of Rab7-positive endosomes (orange arrows). **C–F.** In contrast, in cells depleted of Vps34 (C), Uvrag (D), Vps15 (E), or Atg6 (F), Vps41-9xHA signal becomes largely cytosolic and dispersed. Notably, the large Rab7-positive structures present in these cells are not positive for Vps41-9xHA, indicating defective HOPS recruitment to late endosomal compartments.

**Supplementary Figure 4.**
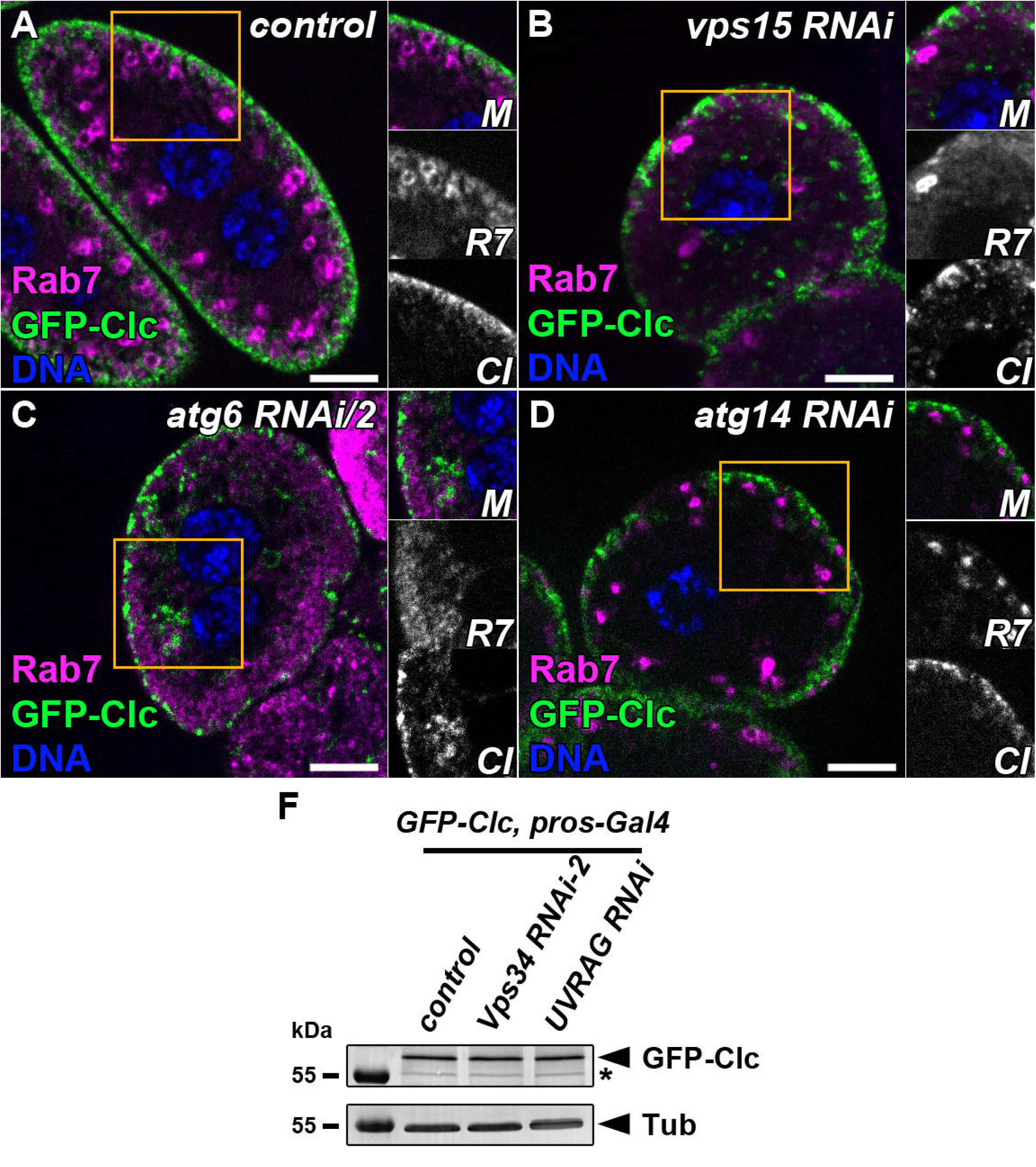
Additional analysis of clathrin localization in PI3K(III) complex mutants. **A–D.** Nephrocytes expressing GFP-Clc were stained with anti-Rab7. Similar to the phenotypes shown in Figure 5, depletion of Vps15 (B) or Atg6 (C) results in the appearance of large GFP-Clc–positive structures deeper in the cytoplasm, some of which colocalize with Rab7. In contrast, Atg14 knockdown cells (D) display a control-like GFP-Clc distribution. Nuclei are shown in blue (DNA). Scale bars: 10 μm. Insets display merged (M), Rab7 (R7), and GFP-Clc (Cl) signals of the boxed regions. **E.** Western blot analysis shows comparable GFP-Clc protein levels in lysates from control, Vps34 RNAi, and Uvrag RNAi larvae, indicating that the clathrin accumulation observed in microscopy images is not due to altered GFP-Clc expression or impaired degradation. *: non-specific band. Tubulin was used as loading control.

**Supplementary Figure 5.**
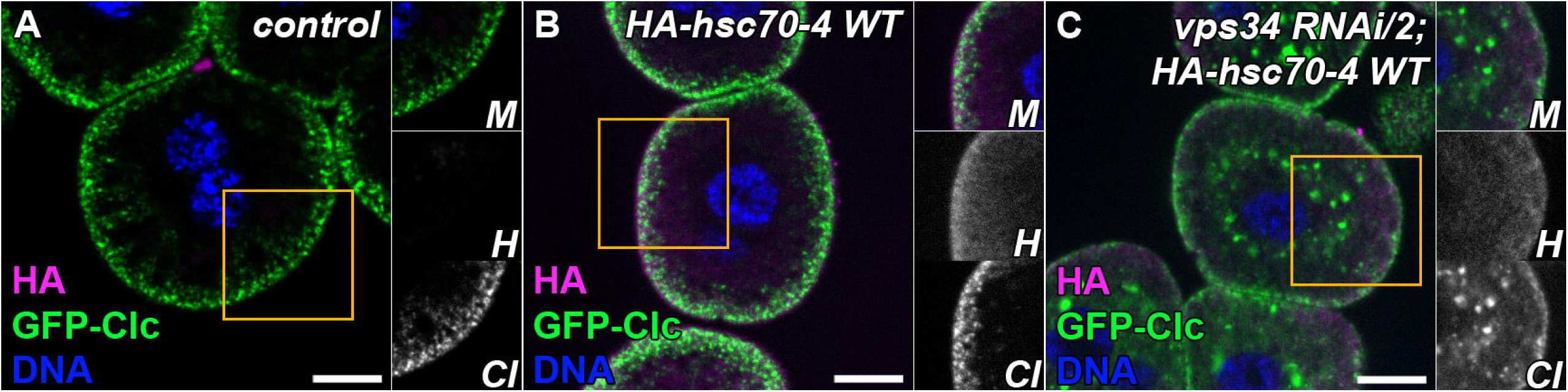
Hsc70-4 partially colocalizes with clathrin-positive structures and becomes more diffuse upon Vps34 depletion. **A–C**. Nephrocytes expressing GFP-Clc were stained with anti-HA antibodies. In blind control cells lacking Hsc70-4–HA expression (A), no HA signal is detected. In Hsc70-4–HA–expressing cells (B), HA signal is predominantly cytosolic but shows enrichment at the cell periphery, where GFP-Clc–positive clathrin-coated endocytic structures are located. In Vps34 RNAi cells (C), this peripheral enrichment is reduced, and Hsc70-4–HA signal appears more uniformly distributed throughout the cytosol. Nuclei are shown in blue (DNA). Scale bars: 10 μm. Insets display merged (M), Hsc70-4-HA (H), and GFP-Clc (Cl) signals of the boxed regions.

